# Iron toxicity potentiates cell-type specific amyloid beta proteotoxicity in *C. elegans* via altered energy homeostasis

**DOI:** 10.64898/2026.03.25.714217

**Authors:** Wilson Peng, Kaitlin B. Chung, Ali Al-Qazzaz, Aidan Straut, M. Kerry O’Banion, B. Paige Lawrence, Robert T. Dirksen, John O. Onukwufor

**Affiliations:** Department of Pharmacology and Physiology, University of Rochester School of Medicine and Dentistry, Rochester NY, 14642 USA; Department of Environmental Medicine and Public Health Sciences, University of Rochester School of Medicine and Dentistry, Rochester NY, 14642 USA; Department of Neuroscience, University of Rochester School of Medicine and Dentistry, Rochester NY, 14642 USA; Del Monte Institute for Neuroscience, University of Rochester School of Medicine and Dentistry, Rochester NY, 14642 USA

**Keywords:** Temperature, Iron Burden, Iron Sensitivity, Tissue-specific Aβ 42 Toxicity, Mitochondrial Dysfunction

## Abstract

Alzheimer’s disease (AD) is a devastating neurodegenerative disorder characterized by memory loss and a decline in cognitive function. Hallmarks of AD include an age-dependent accumulation of toxic amyloid beta (Aβ) 42 in the brain, energy dyshomeostasis caused by mitochondrial dysfunction, and iron overload. However, the role of iron overload and mitochondrial dysfunction in AD pathology is unknown and their precise relationship with Aβ 42 toxicity in AD pathology is unclear. *C. elegans* provide a powerful model system to untangle and clarify these relationships. In this study, we quantify the temperature-dependence of iron toxicity (16, 20 and 25⁰C) in neurons and muscle of *C. elegans* that overexpress Aβ 42. We found that Aβ 42, regardless of the cell-type expression, caused accelerated paralysis compared to age–matched WT worms with the greatest degree of paralysis observed at an elevated temperature (25⁰C). Moreover, the combination of iron toxicity and Aβ 42 results in an enhanced paralytic phenotype at 16⁰C. Thus, iron exposure potentiates Aβ toxicity observed at low temperatures. Iron toxicity stimulated both maximum (State 3) and leak (State 4) respiration in WT and Aβ 42 worms. Aβ 42 worms also exhibited increased leak respiration at baseline that was further exacerbated by iron toxicity. Iron burden and sensitivity increased Aβ 42 peptide toxicity. Aβ 42 worms exhibited reduced levels of Ca, Zn, Mn, and K. Overall, our results suggest that iron potentiates Aβ toxicity at low temperature and enhances Aβ peptide mediated mitochondrial bioenergetic dysfunction in *C. elegans*.

**Graphical Abstract:** 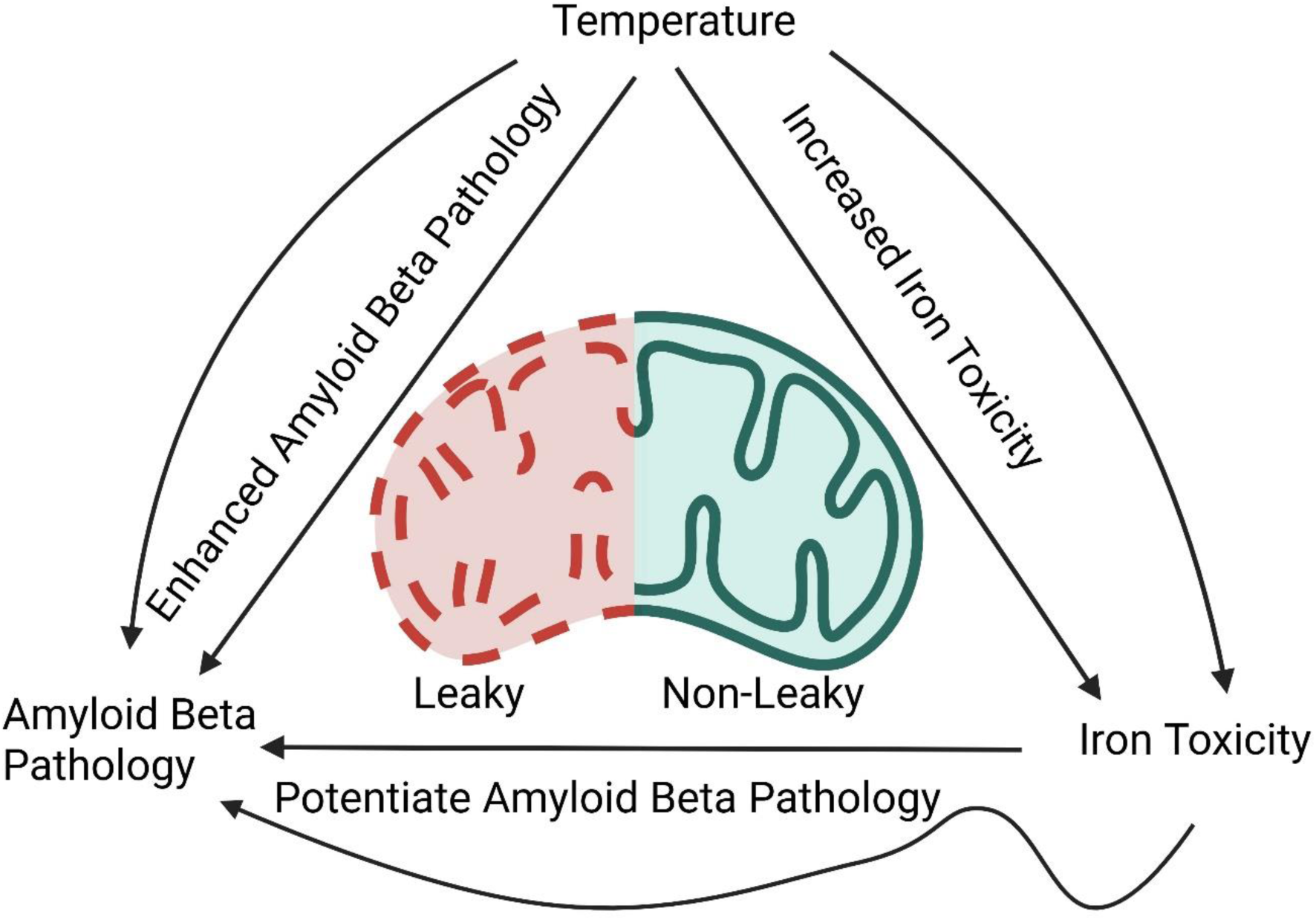

**Highlights:**

1. Temperature stress modulates the synergetic interactions of iron toxicity and Aβ 42 pathology
2. Iron sensitivity drives increased cell-type specific Aβ 42 pathology
3. Energy dyshomeostasis via impaired mitochondrial function and increased proton leak contributes to iron- and Aβ-induced pathology

## 1. Introduction

Alzheimer’s disease (AD) is a devastating chronic progressive neurodegenerative disease and the most common cause of dementia, currently affecting 40 million people world-wide and projected to reach 100 million by the year 2050 [1–3]. AD is characterized by a decline in memory and cognition that worsens with age [4–7]. A major hallmark of AD is the development of amyloid beta (Aβ) pathology, which is thought to drive neuronal loss and death [8, 9]. Aβ peptide is derived from the cleavage of amyloid precursor protein (APP), a type 1 membrane glycoprotein highly expressed in the brain that is encoded by the APP gene on human chromosome 21 [10–12]. Under physiological conditions, APP contributes to processes involved in neuronal homeostasis such as development, signaling, and transport [13–16]. However, under pathological conditions, abnormal cleavage of APP results in the production of toxic levels of Aβ 42 peptide that is thought to promote neuronal AD pathology [5, 17].

Aβ 42 is a peptide that consists of 42 amino acids produced by cleavage of APP by both the β and γ secretases [18]. Abnormal cleavage and/or mutation in genes that regulate β and γ secretase activity results in disproportional production of Aβ 42 relative to the less toxic Aβ 40 isoform [18–21]. Increased levels of toxic Aβ 42 are suggested to contribute to the development and/or progression of AD pathology [18–21]. Consistent with this idea, prior studies demonstrated Aβ 42 toxicity in cell lines [22–25], *C. elegans* [26–31], and mice [32–36]. However, recent reports indicate that clearance and/or reduction of Aβ toxicity may not be sufficient alone to mitigate AD pathology [37–39]. Thus, AD pathology could result from genetic predisposition (e.g., APP mutation resulting in increased Aβ 42 production) combined with environmental stressors such as heavy metal toxicity [40–44].

Iron is a heavy metal that is critical for neuronal health and function such as neurotransmitter production, synaptic activity, metabolism, and serving as a critical co-factor for enzyme activity [45–48]. On the one hand, iron regulates APP expression through activation of iron response elements (IREs) present in the APP promoter [49]. In addition, iron also plays an important role in APP function, including cellular iron export [50]. Specifically, APP possesses iron binding capability through its ferroxidase domain which facilitates the export of iron outside the cell by ferroportin following oxidization of ferrous iron to ferric iron [50–52]. In contrast, others reported no evidence of altered ferroxidase activity by APP [53]. However, APP was recently shown to promote ferrous iron export by stabilizing the cell surface expression of ferroportin [54, 55]. Therefore, this symbiotic relationship between iron and APP promotes neuronal health and function. However, this delicate balance can be disrupted under conditions of iron overload, dysregulation, and/or genetic mutations in APP, ultimately leading to iron- and Aβ 42-mediated neuronal toxicity in AD. Indeed, prior studies found that iron-induced alterations in APP cleavage result in toxic levels of Aβ 42 [50, 56–60]. However, the precise relationship between Aβ 42 and iron toxicity in AD pathology and progression remains unclear.

*C. elegans* provide an excellent model system to address this gap in knowledge and increase understanding of the relationship between iron induced toxicity and Aβ 42 peptide pathology. *C. elegans* with overexpressed human Aβ 42 is a model of AD [26]. In this AD model, worm paralysis, or an inability to move upon stimulation, is observed with Aβ 42 expression and this toxicity is enhanced at elevated temperatures [26–31]. To better understand the relationship between iron toxicity and Aβ 42 pathology, we used wild-type, and Aβ 42 expressed in two different cell-types, neuron and muscle, to determine the tissue specificity/sensitivity of iron-enhanced Aβ 42 toxicity. These studies address the following questions: 1) Are there cell-type-specific effects of the temperature sensitivity of Aβ 42 toxicity (e.g., paralysis)? 2) How does iron modulate tissue-dependent Aβ 42 toxicity? 3) Is iron enhanced Aβ 42 pathology due to increased iron overload and/or sensitivity? 4) What impact does iron burden have on other metals? 5) What is the relationship between iron toxicity, Aβ 42 pathology, and mitochondrial function?

## 2. Materials and Methods

### C. elegans strains and maintenance

The following worm strains were used in this study: N2 (wild type), Roller (worm with circular movement) [61] CL2179 (dvIs179 [myo-3p::GFP::3’ UTR(long) + rol-6(su1006)]), worms carrying human Aβ 42 peptide expressed in muscle including CL4176 (dvIs27 [myo-3p::A-Beta (1-42)::let-851 3’UTR) + rol-6(su1006)] X) and CL2120 (dvls14 [(pCL12) unc-54::Abeta 1-42 + (pCL26) mtl-2::GFP], as well as neurons from CL2355 (dvIs50 [pCL45 (snb-1::Abeta 1-42::3’ UTR(long) + mtl-2::GFP] I). Worms were maintained on OP50 bacterial lawns on nematode growth media (NGM). All strains were maintained at 16⁰C except for N2 which was maintained at 20⁰C. All strains used in this investigation were provided through the *Caenorhabditis* Genetics Center (CGC).

### Thermal sensitivity analysis

Synchronized adult worms from each strain were transferred from their maintenance temperature to an experimental temperature (16, 20 and 25⁰C) either with or without iron (0 and 35 µM) exposure for 10 days. N2 worms were maintained at 20⁰C with measurements performed at 16, 20 or 25⁰C. All other strains (CL2179, CL4176, CL2120 and CL2355) were maintained at 16⁰C, with measurements conducted at 16, 20 and 25⁰C. The concentration of iron (35 µM) used in this investigation was based on our earlier study [31].

### Worm paralysis measurement

Worm paralysis was scored as previously described [31]. In brief, synchronized adult worms were transferred to plate with or without iron (0 and 35 µM) at the desired temperature and experimental conditions. Paralysis, which was defined as the inability to move upon mechanical stimulation, was scored every 24 h at room temperature for the duration of experiment.

### Worm swimming rate measurement

Swimming rate of non-paralyzed worms was measured as previously described [31]. In brief, synchronized adult worms from strains N2, CL4176, CL2179, and CL2355 were transferred to a seeded plate with and without iron (0 or 35 µM) every 24 h until the end of 3 days of exposure period at 25⁰C. Non-paralyzed worms were then transferred individually to an unseeded plate containing 100µl of M9 media (22 mM KH_2_PO_4_, 42 mM Na_2_HPO_4_, 86 mM NaCl, 1 mM MgSO_4_, pH 7). After an acclimatization period of 30 s at room temperature, swimming rate (counts/min) was calculated over 15 s.

### Worm pharyngeal pumping rate measurement

Worm pharyngeal pumping rate was measured with slight modification of an approach described previously [29]. In brief, synchronized adult worms (N2, CL2355, CL4176 and CL2179) were transferred to seeded plates with or without iron (0 and 35 µM) at 25⁰C and then monitored for 3 days. Non-paralyzed worm pharyngeal pumping rate (counts/min) was calculated over 15 s at room temperature.

### Mitochondrial isolation

Worms were first synchronized L1 to adult on HB101 (bacterial lawns), at their cultured temperatures: N2 at 20⁰C, roller and Aβ (neuron and muscle) at 16⁰C. Approximately 0.4 million adult worms were transferred to 25⁰C on fresh seeded HB101 plates with or without iron (0 and 35 µM) for 3 days. To isolate mitochondria, worms were rinsed with M9 media followed by mitochondrial isolation media (225 mM Sucrose, 75 mM Mannitol, 5 mM HEPES, pH 7.3). Using an ice-cold mortar with pure sea sand, worms were crushed, followed by a 60 s homogenization with a handheld motorized homogenizer (Fisherbrand^TM^ Homogenizer 150, Fisher Scientific). After homogenization, the homogenate was centrifuged at a low spin of 800 *g* for 10 min at 4⁰C. The supernatant was then collected and spun at a high speed of 7000 *g* for 10 min at 4⁰C to obtain mitochondrial pellets. The mitochondrial pellets were subsequently washed and spun at a high speed of 7000 *g* for 5 min at 4⁰C. The mitochondrial pellets were resuspended in an equal volume of isolation media with mitochondrial protein determined by the Folin-phenol method.

### Mitochondrial respiration measurement

Mitochondrial respiration measures were obtained using a Clark-type O_2_ electrode (OxyGraph^+^, Hansatech Instruments) as described previously [31]. In brief, following calibration of the chamber at 25⁰C, freshly isolated mitochondria (1 mg/ml) were added to the chamber. To spike the TCA cycle, complex-1 linked substrates (2.5 mM malate and 5 mM glutamate) were added to generate complex-1 linked reducing equivalent. The addition of 0.4 mM ADP resulted in the stimulation of maximum respiration (state 3 respiration). Then subsequent addition of 1 µg/ml of oligomycin to inhibit complex V enabled quantification of leak respiration (state 4 respiration). The respiratory control ratio (RCR) was calculated from the ratio of state 3 respiration over that of state 4 respiration. In all conditions, substrates were added to the chamber via a syringe port.

### Whole worm metal burden

Metal tissue burden in worms was measured using Inductively Coupled Plasma – Mass Spectrometry (ICP-MS) as described previously [31]. In brief, synchronized adult worms (∼20,000) of wildtype, roller, Aβ neuron and Aβ muscle were transferred to a seeded plate with or without iron (0 or 35 µM) and maintained at 25⁰C for 3 days. Worms were washed with media (M9) and then collected by slow centrifugation at 2000 *g* for 2 min. Pellets were resuspended in M9 to a total volume of 1 mL. Worm samples were weighed before digestion. Samples were digested in 1 mL of 69% HNO_3_ and 0.5 mL HCl (36%) at 100⁰C for 60 min. After digestion, samples were cooled, and 10 ml total volume of double deionized water was added to the samples. Samples were analyzed by ICP-MS (Perkin Elmer 2000C) using the following detection limits in ppb (ng/ml): Fe 0.123, Cu 0.0009, Zn 0.0241, Mn 0.00136, Ca 3.688, S 219.3, K 1.706, Mg 0.023, Na 0.088, and P 0.566. Data were normalized to the initial sample weight in grams.

### Temperature coefficient (Q_10_ values) calculation

Temperature sensitivity was calculated as previously described [62]. In brief, temperature coefficient values (Q_10_) for paralysis of wild type worms (N2), roller worms, and worms with Aβ expressed in either muscle or neuron were calculated for temperature ranges of 16-20⁰C, 20-25⁰C and 16-25⁰C using the van’t Hoff equation:

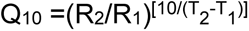

Where R_2_ and R_1_ represent worm paralysis at two temperatures T_2_ and T_1_ and where T_2_>T_1_. We used mean values of the percent paralyzed worms to calculate Q_10_ for both iron- and/or temperature-exposed worms.

### Statistical analyses

All data were subjected to normality and homogeneity variance testing, and all data passed the test. Data were analyzed using two-way ANOVA with Tukey post hoc test for multiple comparisons. Results were considered significant at p-value < 0.05. Statistical analyses were conducted with GraphPad^TM^ Prism v10 (GraphPad Software, San Diego, CA, USA).

## 3. Results

### 2.1. Iron toxicity enhances cell-type specific amyloid beta thermal sensitivity induced paralysis

Previous studies reported that overexpression of human Aβ peptide in worms enhances toxicity observed at high temperature [63]. To further evaluate the underlying mechanisms involved, we first assessed the effects of temperature and iron toxicity on wild type (WT) worms (Fig. 1A-C). WT worms maintained at 16⁰C showed noticeable paralysis on day 7, with exposure to 35 µM iron shifting this one day earlier (to day 6) and marginally increasing the maximal effect observed at 10 days (Fig. 1A). When WT worms were maintained at 20⁰C, paralysis was detected even earlier during iron exposure (on day 4) and the maximal effect at day 10 was also larger (Fig. 1B). Iron exposure resulted in a further acceleration in the rate (to day 3) and degree of paralysis with an additional 5⁰C increase in temperature (from 20 to 25⁰C) (Fig. 1C). Thus, iron exposure significantly enhanced both the onset and magnitude of temperature-dependent paralysis of WT worms.

**Fig. 1:**
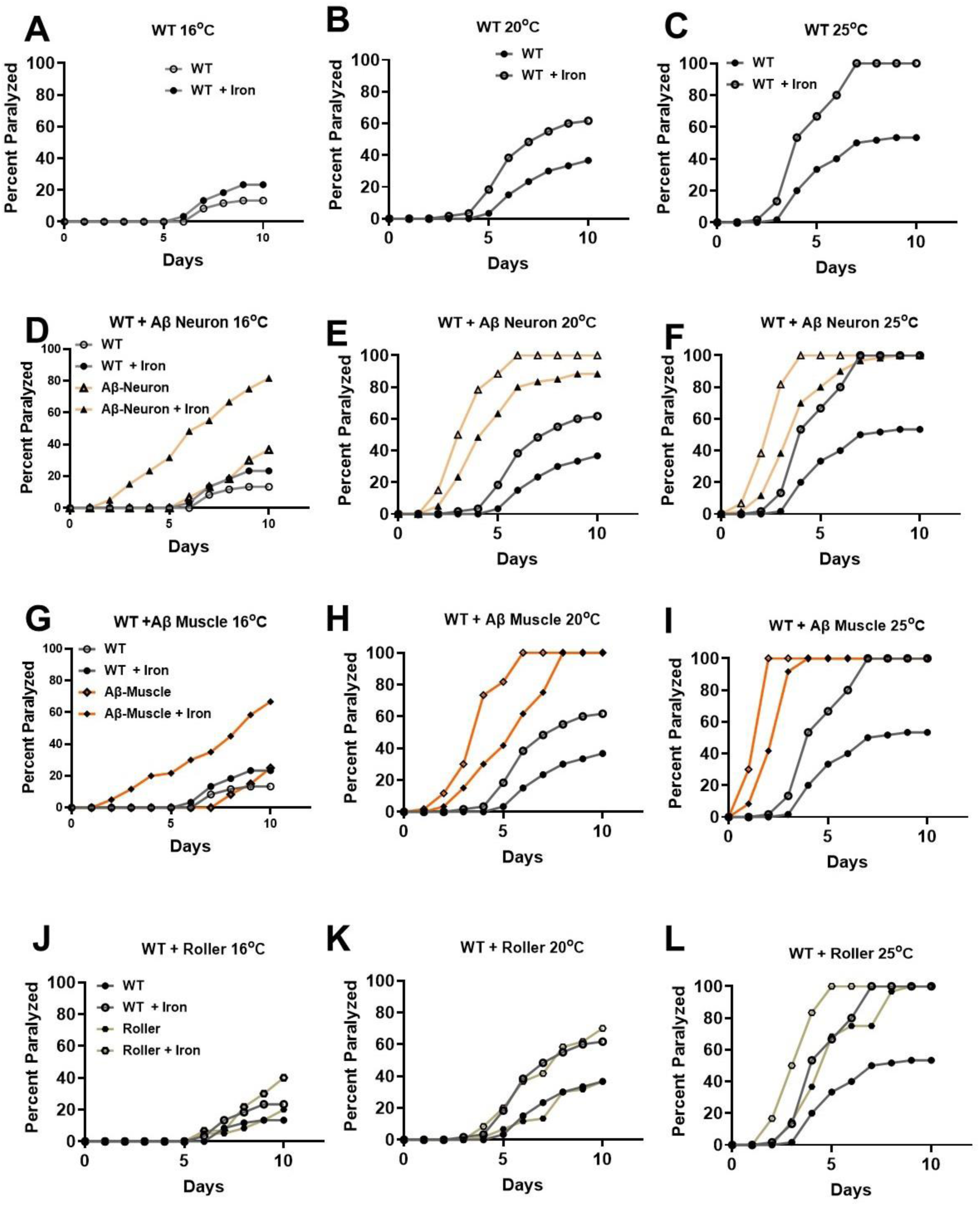
Iron toxicity potentiates Aβ 42 peptide thermal sensitivity. Synchronized L4 worms from wild-type WT (A-C), Aβ neuron (D-F), Aβ muscle (G-I), and roller (J-L) were transferred to a plate containing iron (0 and 35 µM) and assigned to temperatures (16, 20 and 25⁰C). Worms were transferred every 24h for 10 days. Worms were scored for paralysis (e.g., inability to move upon stimulation) every 24h for 10 days. **Data from WT was embedded in the rest of the treatments to see their deviations from WT**. Data are mean of N=3 independent biological replicate (where one biological replicate contains 20 worms per plate).

Neurons are reported to be sensitive to Aβ 42 peptide toxicity [30, 64]. However, the specific impact of neuronal Aβ 42 peptide expression on the iron-induced enhancement of temperature-dependent worm paralysis is unknown. Worms with overexpression of Aβ 42 peptide are typically cultured at 16⁰C to limit Aβ 42 expression and toxicity [26]. When we assessed worms with neuronal-specific Aβ 42 peptide maintained at 16⁰C, we first observed paralyzed worms on day 6 (similar to WT worms) in the absence of iron and as early as day 2 in the presence of iron (Fig. 1D). Thus, neuronal Aβ 42 peptide expression resulted in a marked (4-day) shift in the onset of paralyses, consistent with neuronal Aβ 42 peptide potentiating iron toxicity. In total, about 30% and 75% worms were paralyzed on day 10 at 16⁰C in the absence and presence of iron exposure, respectively (Fig. 1D). When the temperature was increased to 20⁰C, the onset of paralysis occurred on day 2 both with and without iron exposure, though the magnitude of paralysis was greater in worms exposed to iron (Fig. 1E). In contrast, when neuronal Aβ 42 worms were maintained at 25⁰C, all worms were paralyzed by day 10, though iron exposure accelerated the time-dependence of this paralysis (Fig. 1F). Overall, iron exposure and neuronal expression of Aβ 42 peptide synergistically potentiated temperature-dependent worm paralysis.

A similar analysis of an interaction between iron and Aβ 42 peptide toxicity in the temperature-dependence of paralysis was assessed for worms with muscle-specific Aβ 42 peptide expression. For worms with muscle-specific Aβ 42 expression maintained at 16⁰C, the onset of paralysis was not until day 8 (Fig. 1G), which was the longest delay in paralysis observed for all conditions evaluated in Fig. 1. These findings suggest that expression of Aβ 42 peptide may provide a modest level of protection against temperature-dependent worm paralysis in the absence of iron. However, iron exposure of muscle-specific Aβ 42 worms maintained at 16⁰C produced a dramatic increase in both the onset of paralysis (from day 8 to day 2) and increase in percent of worms paralyzed (from 20% to 70%) on day 10 (Fig. 1G). When the temperature was increased to 20⁰C, the onset of paralysis without iron occurred earlier (day 2) compared to that observed at 16°C (day 8, Fig. 1G) or WT worms at 20⁰C (day 5, Fig. 1B), suggesting a synergistic effect between temperature and muscle-specific Aβ 42 peptide expression. By day 10, all control and iron-treated muscle-specific Aβ 42 peptide worms were paralyzed (Fig. 1H) when maintained at either 20⁰C (Fig. 1H) or 25⁰C (Fig. 1I), although iron exposure accelerated the time-dependence of this paralysis. Thus, as observed for neuronal Aβ 42 worms, iron exposure similarly potentiated temperature-dependent worm paralysis of muscle-specific Aβ 42 expressing worms (Fig. 1G-I). However, nearly all iron-treated, muscle-specific Aβ 42 expressing worms maintained at 25⁰C were paralyzed by day 2 (Fig. 1I), while only ∼40% of iron-treated neuron-specific Aβ 42 expressing worms were paralyzed by day 2 (Fig. 1F).

Since muscle-specific Aβ 42 peptide expressing worms exhibited a roller phenotype (circular movement) [61], there is the possibility that the roller phenotype underlies the more profound paralysis observed in these worms. To test this idea, we quantified temperature/iron-dependent paralysis in a line of worms with a roller phenotype that lack Aβ 42 peptide expression (Fig. 1J-L). The delayed onset of paralysis observed at 16⁰C in muscle-specific Aβ 42 peptide expressing worms was not replicated in roller worms lacking Aβ 42 peptide expression (Fig. 1J vs 1G), suggesting that the delayed paralysis onset requires muscle Aβ 42 peptide expression. Furthermore, the prominent early-onset paralysis documented for muscle-specific Aβ 42 peptide worms treated with iron at 16⁰C (Fig. 1G) was absent in roller worms exposed to iron at 16⁰C (Fig. 1J). In addition, while muscle-specific Aβ 42 peptide worms were >90% paralyzed by day 3 at 25⁰C (Fig. 1I), only ∼10% of roller worms were paralyzed on day 3 at 25⁰C (Fig. 1L). Indeed, the temperature dependence of paralysis for roller worms in both the absence and presence of iron (Fig. 1J-L) were similar to that of WT worms (Fig. 1A-C). To further evaluate the impact of muscle specific Aβ peptide expression, we assessed the effect of using a different promoter to drive the expression of Aβ peptide in muscle. Since worms with muscle-specific Aβ peptide expression with a roller phenotype used the *myo-3* promoter, we compared this with muscle-specific Aβ peptide expression using the *unc-54* promoter, which lacked a roller phenotype. The delayed paralysis observed at 16⁰C with *myo-3* promoter was absent with the *unc-54* promoter (S Fig. 1A) and also lacked iron-induced enhancement of Aβ 42 peptide toxicity (S Fig. 1A). In addition, the enhanced temperature dependence of paralysis observed with *myo-3* promoter was also absent with the *unc-54* promoter (S Fig. 1B&C). Overall, the impact of Aβ 42 peptide expression in muscle on worm paralysis primarily resulted from interactions between temperature, iron exposure, and the promoter driving muscle-specific Aβ expression rather than a specific function of their roller phenotype.

### 2.2. Effects of iron and tissue-specific Aβ 42 expression on the temperature-dependence of paralysis

Temperature is a critical factor that regulates a wide range of cellular functions and stress susceptibility [62, 65]. The temperature coefficient of a process (or Q_10_-value) assesses the relative degree of responsiveness of the process to a change in temperature [62]. To determine how different cell type-specific expression of Aβ 42 peptide impacts the temperature dependence of paralysis, we calculated Q_10_-values for each genotype and treatment condition studied in Fig. 1 (S Table1). Since Aβ worms were all paralyzed at 25⁰C, analysis of Q_10_ value was limited to the 16-20⁰C temperature range. The Q_10_ value for WT worms in the absence of iron with a temperature change from 16 to 20⁰C was 8.69, which was even larger (11.36) in the presence of iron, consistent with iron increasing the temperature response of WT worms. These results suggest that iron exerts a modest increase in the temperature dependence of WT worm paralysis. Neuronal expression of Aβ 42 peptide exhibited a minimal increase in the Q_10_ value of WT worms from 16 to 20⁰C in the absence of iron (9.01 vs 8.69). Interestingly, iron exposure drastically reduced the Q_10_ value in this temperature range to 1.66, which was far lower than that of WT worms (11.36). In contrast, muscle-specific Aβ 42 peptide expression exhibited the greatest temperature responsiveness for the 16-20⁰C temperature range with a Q_10_ value of 32.00 in the absence of iron. These findings suggest that temperature exerts a larger effect on Aβ 42 peptide toxicity in muscle compared to that observed in WT and neuronal Aβ 42 worms. In the presence of iron, a Q_10_ value of 2.76 was observed for muscle-specific Aβ 42 worms for the temperature range of 16-20⁰C. Because muscle Aβ 42 peptide worms are rollers, we also assessed the effect of the roller phenotype on Q_10_ values. At the temperature range of 16-20⁰C, rollers had Q_10_ values of 4.55 and 4.05 in the absence and presence of iron, respectively. Similarly, muscle Aβ 42 with *unc-54* promoter had Q_10_ values of 3.74 in the absence of iron and 4.70 in the presence of iron, suggesting a marginal increase with iron exposure like WT. Thus, the high Q_10_ value (32.00) observed for muscle-specific Aβ 42 in this temperature range depended on muscle-specific Aβ 42 peptide together with the promoter driving the expression and not the roller phenotype.

### 2.3. Cell type-specific impact of amyloid beta peptide and iron toxicity on mitochondrial function

Our data indicate that cell type specific Aβ 42 peptide toxicity is enhanced with increasing temperature (Fig. 1). Furthermore, the temperature responsiveness (Q_10_) of Aβ 42 peptide toxicity depends on the tissue (muscle vs neurons) where Aβ 42 peptide is expressed. To more critically examine the cell type specificity of Aβ 42 peptide toxicity and the impact of iron on this toxicity, we assessed paralysis, swimming rates of non-paralyzed worms, and pumping rates at 25⁰C for 3 days. We found that Aβ 42 peptide expression enhanced worm paralysis regardless of the cell type where Aβ 42 peptide was expressed (i.e., muscle or neurons) (Fig. 2A). However, in the absence of iron exposure, worms with muscle-specific Aβ 42 peptide expression exhibited greater paralysis than neuron-specific Aβ 42 peptide expressing worms (Fig. 2A, *left*). Moreover, while iron exposure worsened paralysis across all genotypes (WT, roller, Aβ-neurons, Aβ-muscle), iron exposure produced the greatest degree of paralysis in muscle-specific Aβ 42 peptide expressing worms (Fig. 2A, *right*). However, after adjusting for the roller phenotype, Aβ muscle and neuron paralysis was similar at baseline with neuron-specific Aβ 42 worms exhibiting greater paralysis with iron exposure than muscle-specific Aβ 42 worms (S Fig. 2A).

**Fig. 2:**
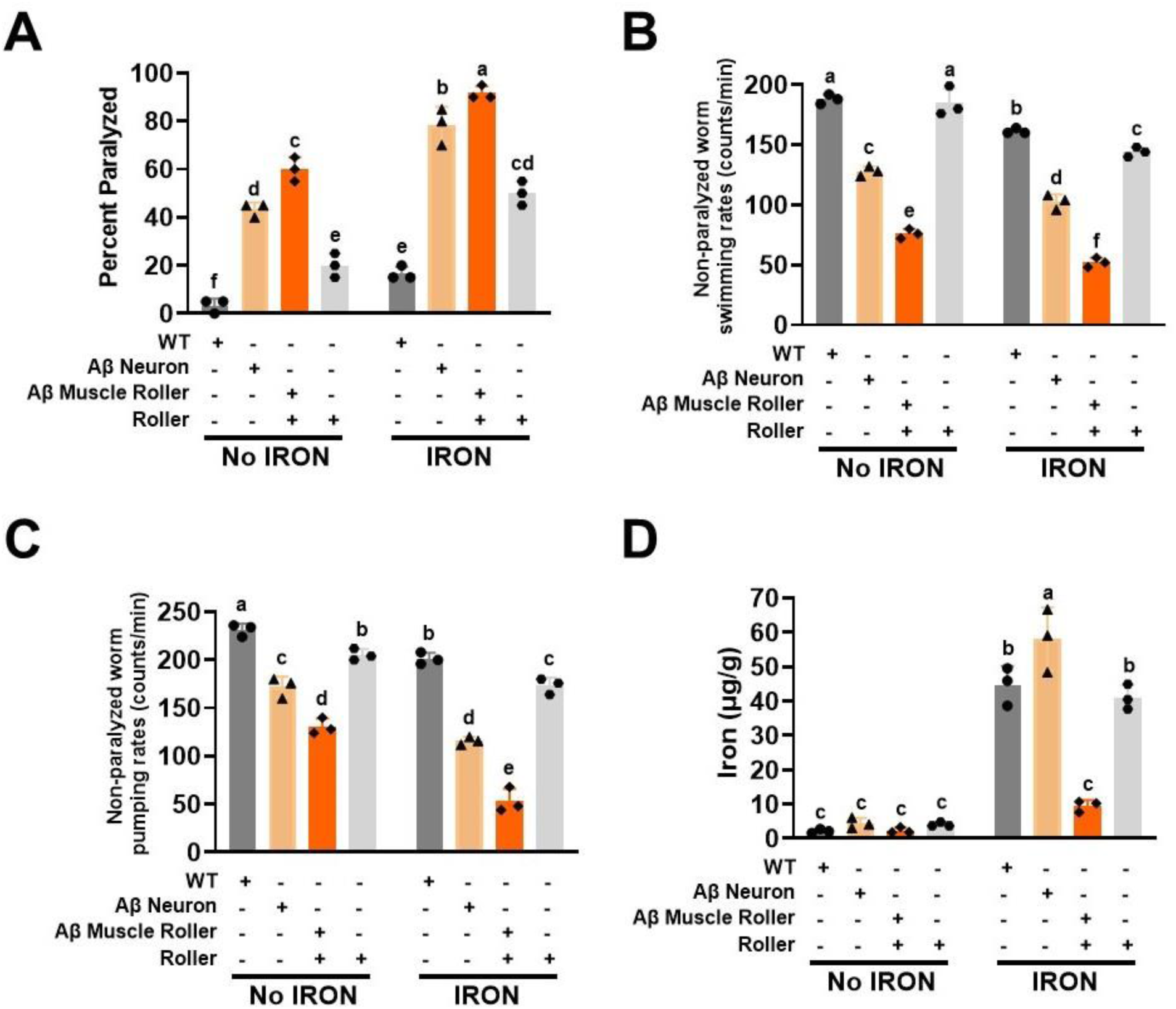
Impact of iron toxicity on Aβ 42 peptide is cell-type specific. A) Iron worsens Aβ 42 peptide induced paralysis. Synchronized L4 worms from WT, roller, Aβ neuron, and Aβ muscle genotypes were transferred to a plate containing iron (0 and 35 µM) and incubated at 25⁰C for three days. Paralysis was scored for 3 days. Data are mean ± SEM N= 3 independent biological replicates (where one biological replicate contains 20 worms per plate). Bars with different letters differ significantly, two-way ANOVA, Tukey post hoc test. B) Iron toxicity exacerbates Aβ 42 peptide effects on swimming rate. Synchronized L4 worms from WT, roller, Aβ neuron, and Aβ muscle genotypes were transferred to a plate containing iron (0 and 35 µM) and incubated at 25⁰C for three days. Non-paralyzed worms were individually transferred to an unseeded plate containing 100 µl of buffer. A 30s conditioning period was observed, swimming rates were collected for 15s. Data are mean ± SEM, N = 3 independent replicates (where 5 independent worm count constitute an N). Bars with different letter differs significantly, two-way ANOVA, Tukey post hoc test. C) Iron toxicity worsens Aβ 42 peptide effects on pumping rate. Synchronized L4 worms from WT, roller, Aβ neuron, and Aβ muscle genotypes were transferred to a plate containing iron (0 and 35 µM) and incubated at 25⁰C for three days. Worm pumping rates were collected for 15s. Data are mean ± SEM, N = 3 independent replicates (where 5 independent worm count constitute an N). Bars with different letter differs significantly, two-way ANOVA, Tukey post hoc test. D) Increased tissue iron burden is not responsible for Aβ 42 pathology. Synchronized L4 worms (20,000 worms) from WT, roller, Aβ neuron, and Aβ muscle genotypes were transferred to a plate containing iron (0 and 35 µM) and incubated at 25⁰C for three days. Iron burden was quantified by ICP-MS. Data are mean ± SEM, N=3 independent biological replicates (where one biological replicate contains 20,000 worms per plate). Bars with different letter differs significantly, two-way ANOVA, Tukey post hoc test.

We previously reported that swimming rates of non-paralyzed worms declines with age and that iron exposure potentiates this effect [31]. Worm swimming rate is a marker of both a decline in energetic output and a progessive loss of function that culminates in paralysis [31]. We confirmed that iron exposure reduced the swimming rate of non-paralyzed WT, roller, and Aβ 42 peptide expressing worms (Fig. 2B, *right*). Moreover, in the absence of iron, swimming rates of both neuronal- and muscle-specific Aβ 42 peptide expressing worms were lower than that of WT and roller worms, with Aβ 42 peptide expression in muscle exhibiting the strongest effect (Fig. 2B, *left*) even after accounting for the roller effect (S Fig. 2B, *left*), which was exacerbated by the presence of iron (S Fig. 2B, *right*).

Because the roller phenotype of muscle specific Aβ worms could have contributed to their slow swimming rates and increase paralysis at baseline, we quantified an alternative functional readout; pharyngeal pumping rate. Worm pharyngeal pumping rate is a proxy measure of worm feeding behavior [66–68]. At baseline (Fig. 2C, *left*), muscle spcific Aβ 42 exhibited the lowest pumping rate with iron further exacebating this effect (Fig. 2C, *right*). Since the pumping rate of rollers was significantly reduced at baseline relative to WT worms (Fig. 2C, *left*), we accounted for this by adjusting the roller effect on muscle specific Aβ 42 (S Fig. 2C, left). The adjusted values revealed that both Aβ muscle and neuron worms exhibited similar pumping rates at baseline (S Fig. 2C, *left*). However, in the presence of iron, Aβ muscle worms exhibited a lower pumping rate than that of Aβ neuron worms (S Fig. 2C, *right*).

Since iron exposure exacerbated the effect of Aβ 42 peptide expression in neurons and muscle on paralysis, swimming rate and pumping rate (Fig. 2A-C), we tested if Aβ 42 peptide genotypes exhibited a higher overall iron burden relative to WT worms. As expected, iron burden was low and not significantly different across all four genotypes in the absence of iron exposure (Fig. 2D, *left*). However, following iron exposure, WT, roller, and neuronal Aβ 42 peptide exhibited a higher level of iron burden compared to muscle Aβ 42 peptide worms, which exhibtied the lowest iron burden (Fig. 2D, *right*). These results suggest that the impact of iron on Aβ 42 peptide toxicity in muscle was due to increased iron sensitivity rather than iron overload. We hypothesized that the slower pumping and swimming rates, and increased paralysis of Aβ 42 expressing worms resulted from reduced levels of and/or dysfucntional mitochondria. To test this hypothesis, we isolated mitochondria from worms of all four genotypes under the same iron and iron-free conditions in which pumping rate, swimming rate, and paralysis were measured (Fig. 2A-C). At baseline, maximium respiration (state 3) was lower for the Aβ muscle and roller genotypes (Fig. 3A, *left*). Iron exposure stimulated maximum respiration in all genotypes with Aβ neuron exhibiting the highest degree of stimulation (Fig. 3A, *right*) and both roller genotypes exhibiting the lowest level of leak respiration (Fig. 3B, *left*). Exposure to iron enhanced leak respiration across all genotypes, with Aβ neuron worms being the most impacted (Fig. 3B, *right*). Both WT and roller genotypes exhibited a similar mitochondrial respiratory control ratio (RCR), a measure of mitochondrial ATP production capacity (cellular energy currency), both at baseline (Fig. 3C, *left*) and after iron exposure (Fig. 3C, *right*). These results suggest that the roller phenotype did not impact mitochondrial function. We observed a significantly reduced RCR in worms with Aβ 42 peptide expression in neurons and muscle (Fig. 3C, *left*). These findings are consistent with the reduction in pumping and swimming rates of Aβ worms being due in part to inefficient mitochondrial function at baseline, with this reduction being further exercebated by iron exposure.

**Fig. 3.**
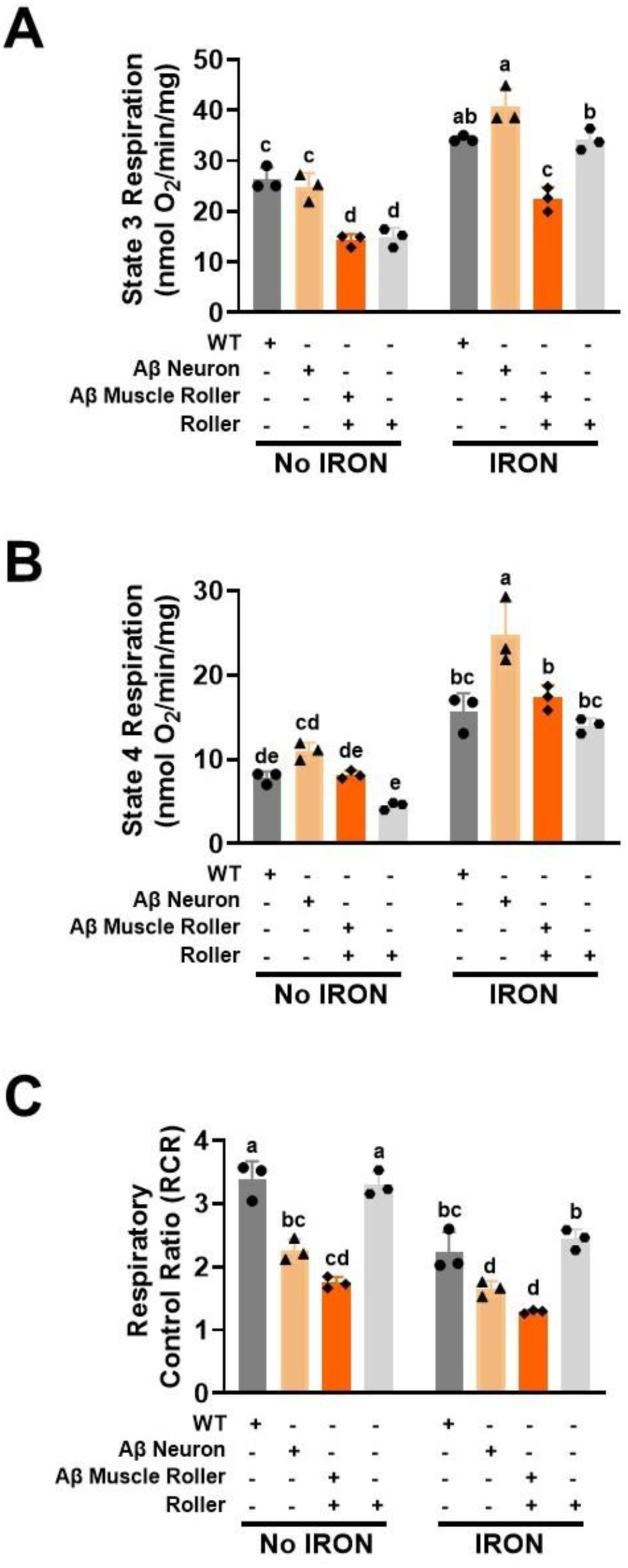
Iron toxicity worsens Aβ 42 peptide mitochondrial dysfunction. Synchronized L4 worms (0.4 million) from WT, roller Aβ neuron and Aβ muscle were transferred to a seeded plates containing iron (0 and 35 µM) at 25⁰C. Mitochondria were isolated after 3 days and mitochondrial respiratory function was quantified. A) State 3 respiration, B) State 4 (Leak) respiration and C) Respiratory control ratio (RCR) maximum/minimum respiration. Data are mean ± SEM N = 3 independent replicates. Bars with different letters differ significantly, two-way ANOVA, Tukey post hoc test.

We conducted correlation analyses to further probe the relationship between iron exposure and cell type specific Aβ 42 peptide expression on pumping rate, swimming rate, paralysis, and iron burden. We observed distinct positive and negative relationships, that were either strong or weak, depending on readout and genotype (Fig. 4). As expected, swimming rate and paralysis exhibited a negative correlation (Fig. 4A) according to the following order: roller (R^2^= 0.72), WT (R^2^= 0.75), Aβ neuron (R^2^= 0.87), and Aβ muscle (R^2^= 0.95). A positive correlation was observed between paralysis and iron burden (Fig. 4B): WT (R^2^= 0.81), Aβ neuron (R^2^= 0.85), roller (R^2^= 0.90) and Aβ muscle (R^2^= 0.90). A similar positive correlation was observed between swimming rate and pumping rate (Fig. 4C): roller (R^2^= 0.64), Aβ neuron (R^2^= 0.82), Aβ muscle (R^2^= 0.89) and WT (R^2^= 0.90). In contrast, a strong negative relationship was observed between pumping rate and paralysis (Fig. 4D): WT (R^2^= 0.77), Aβ muscle (R^2^= 0.89), roller (R^2^= 0.94), and Aβ neuron (R^2^= 0.95). A similar strong negative association was observed between swimming rate and iron burden (Fig. 4E): neuron (R^2^= 0.78), Aβ muscle (R^2^= 0.88), roller (R^2^= 0.90) and WT (R^2^= 0.97). Finally, a strong nagetive correlation was observed between pumping rate and iron burden (Fig. 4F) in the following order: roller (R^2^= 0.82), WT (R^2^= 0.85), Aβ neuron (R^2^= 0.88) and Aβ muscle (R^2^= 0.99). The results in Fig. 4 are consistent with iron burden being responsible for the reduced pumping rate, swimming rate, and increased paralysis observed in cell type specific Aβ 42 peptide expressing worms. In particular, the strong correlations between iron burden and swimming rate (R^2^ = 0.88), paralysis (R^2^ = 0.90), and pumping rate (R^2^ = 0.99), in spite of the relatively low levels of iron burden observed in muscle-specific Aβ 42 peptide expressing worms (∼5-fold lower than WT, roller and neuronal Aβ 42 worms) suggests that the sensitvity to iron toxicity is exceptionally high in these worms. Overall, our data support a differential effect of iron on Aβ 42 toxicity that depends on the cell-type and level of Aβ 42 peptide expression.

**Fig. 4.**
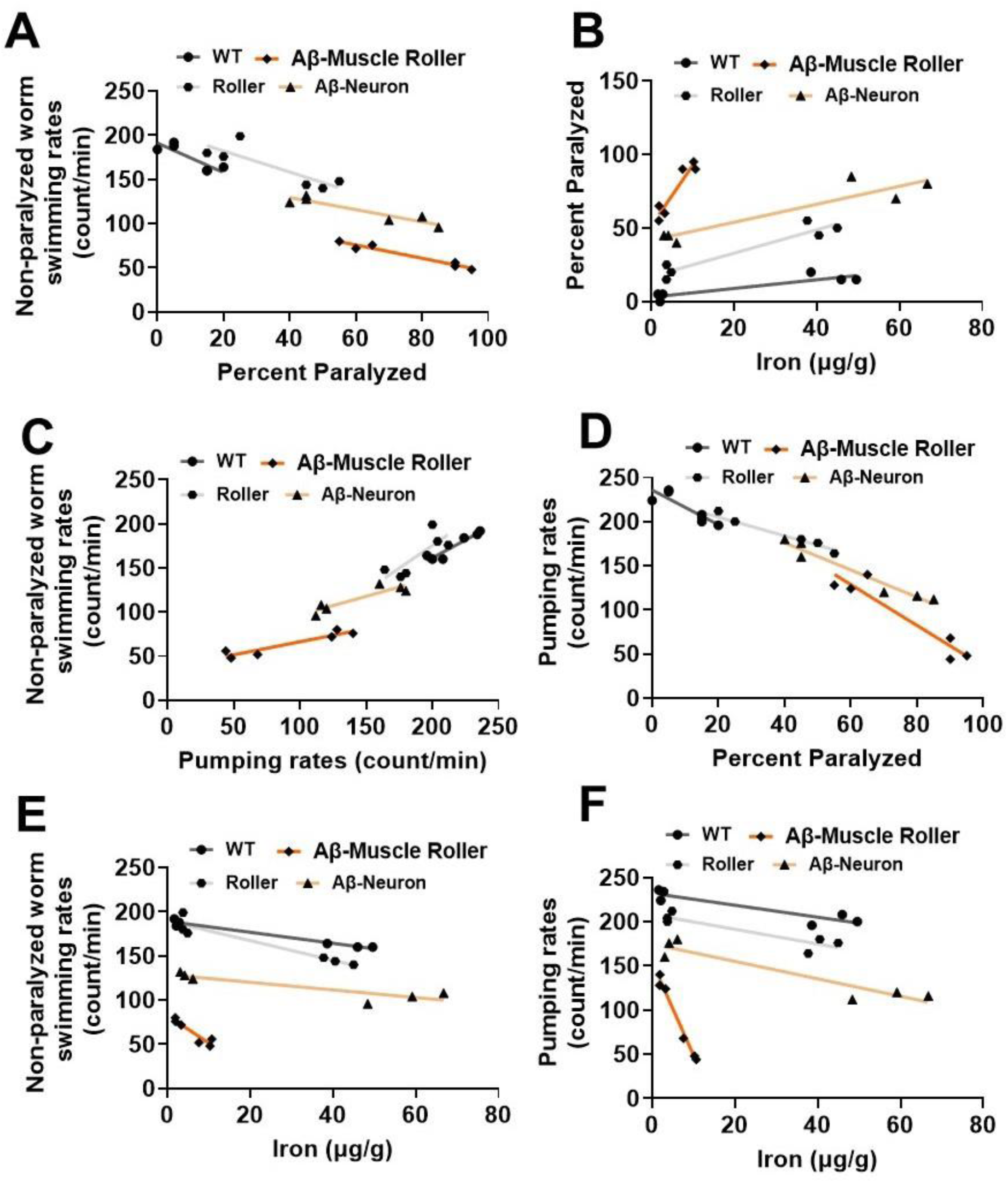
Divergent relationships between iron and Aβ 42 peptide: A). Negative correlation between swimming rate and paralysis in the following order: roller (R^2^= 0.72), WT (R^2^= 0.75), Aβ neuron (R^2^= 0.87) and Aβ muscle (R^2^= 0.95). B). Positive correlation between paralysis and iron burden in the following order: WT (R^2^= 0.81), Aβ neuron (R^2^= 0.85), roller (R^2^= 0.90) and Aβ muscle (R^2^= 0.90). C). Positive correlation between swimming rate and pumping rate in the following order: roller (R^2^= 0.64), Aβ neuron (R^2^= 0.82), Aβ muscle (R^2^= 0.89) and WT (R^2^= 0.90). D). Negative correlation between pumping rate and paralysis in the following order: WT (R^2^= 0.77), Aβ muscle (R^2^= 0.89), roller (R^2^= 0.94), and Aβ neuron (R^2^= 0.95). E). Negative correlation between swimming rate and iron burden in the following order: neuron (R^2^= 0.78), Aβ muscle (R^2^= 0.88), roller (R^2^= 0.90) and WT (R^2^= 0.97). F). Negative correlation between pumping rate and iron burden in the following order: roller (R^2^= 0.82), WT (R^2^= 0.85), Aβ neuron (R^2^= 0.88) and Aβ muscle (R^2^= 0.99). All data are mean from figure 2 (N=6) for each condition.

**Fig. 5:**
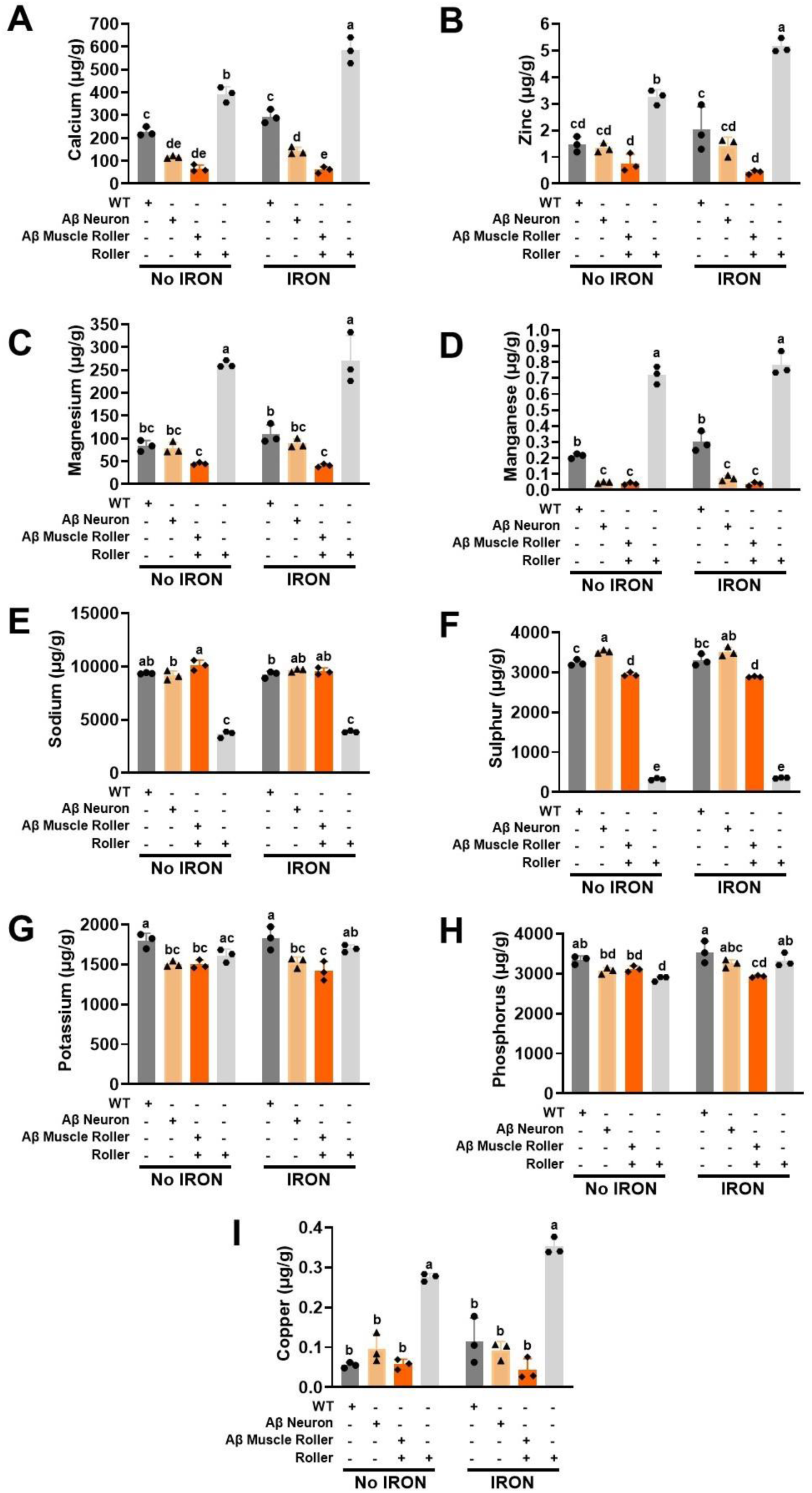
Genotype not Iron burden modulates resident metal burden: Synchronized L4 worms (20,000 worms) from WT, roller, Aβ neuron, and Aβ muscle genotypes were transferred to a plate containing iron (0 and 35 µM) and incubated at 25⁰C for three days. Elemental burden was quantified by ICP-MS. A) Calcium, B) Zinc, C) Magnesium, D) Manganese, E) Potassium, F) Sulfur, G) Sodium, H) Phosphorus, and Copper. WT, roller, Aβ neuron, and Aβ muscle strains were all included in the analysis to compare body burden with and without iron exposure. Data are mean ± SEM, N=3 independent biological replicates (where one biological replicate contains 20,000 worms per plate). Bars with different letters differ significantly, two-way ANOVA, Tukey post hoc test.

### 2.4. Iron toxicity modulates burden of specific metal ions in the context of cell type-specific Aβ 42 peptide expression

Our results revealed that the degree of iron toxicity depends on a combination of iron burden, sensitivity, and cell type-specific Aβ 42 peptide expression (Fig. 2A-D). We tested if an increase in iron uptake and/or sensitivity would disproportionately impact the burden of other resident metals. To test this hypothesis, we quantified levels of nine metal ions (Ca, Zn, Mg, Mn, K, S, Na, P and Cu) across the four genotypes both in the absence and presence of iron exposure (Fig. 4A-I). Iron exposure significantly increased Ca levels in roller, with no effect on WT, neuronal-specific Aβ 42, or muscle-specific Aβ 42 worms (Fig. 4A). Total Ca levels were significantly reduced in Aβ 42 expressing worms, with muscle-specific Aβ 42 worms exhibiting the lowest levels of Ca under both control and iron exposed conditions (Fig. 4A). Iron exposure significantly increased Zn burden in roller, with muscle-specific Aβ 42 worms having the lowest Zn burden (Fig. 4B). Mg burden was not significantly impacted by iron exposure in any of the four genotypes with roller worms exhibiting the highest Mg burden, while muscle-specific Aβ 42 worms exhibiting the lowest Mg burden (Fig. 4C). Iron exposure had no effect on Mn burden in WT, roller, or Aβ 42 expressing, worms (Fig. 4D). In additon, both neuronal- and muscle-specific Aβ 42 worms exhibited a significantly reduced Mn burden compared to roller and WT worms, both in the absence and presence of iron (Fig. 4D). Similar to Mn, K burden was lower in Aβ worms (Fig. 4E), while S (Fig. 4F) and Na (Fig. 4G) were very low in roller worms. P was significantly reduced in iron-exposed, muscle-specific Aβ worms and increased in roller worms (Fig. 4G). No significant changes in Cu burden (Fig. 4I) was observed with iron exposure, though roller worms exhibited an elevated Cu burden. Overall, the results in Fig. 4 indicate that genotype and cell-type specific Aβ 42 expression significantly impacted metal ion burden, but iron exposure did not.

## 4. Discussion

Temperature modulates multiple processes in organismal biology and/or pathophysiology. In worms, the Aβ 42 peptide toxicity is reduced at low temperatures but is enhanced at higher temperatures [63]. Several studies capitalized on this dynamic to probe Aβ 42 peptide toxicity in worms [29–31]. Here, we confirmed that across genotypes, an increase in temperature reduced swimming and pumping rates and enhanced paralysis with a greater impact observed in worms carrying Aβ 42 pathology (Fig. 1). However, the tissue-specific impact of Aβ 42 peptide expression on these phenotypes and how this is modified by iron exposure is unknown. Our results demonstrate tissue-specific Aβ 42 peptide expression enhanced iron toxicity and Aβ 42 peptide-associated pathologies.

The precise relationship (e.g. independent, synergistic) between temperature-, iron-and Aβ 42 peptide-related pathologies in worms is unclear since all three reduce swimming and pumping rates, and increased paralysis. To help untangle this complex relationship, we tested if iron toxicity activates dormant Aβ 42 peptide toxicity. Specifically, we assessed the impact of iron exposure at a temperature (16⁰C) where Aβ 42 peptide induced paralysis was minimal. Consistent with a synergistic interaction, we found that iron exposure enhanced Aβ 42 peptide toxicity (both timing and magnitude) at this temperature. The Aβ 42 peptide toxicity observed at 16⁰C could only be attributed to the effect of iron because this effect was not replicated in either WT or roller worms. Furthermore, we found that the impact of iron toxicity on these Aβ 42 peptide associated pathologies differed depending on where Aβ 42 peptide was expressed (i.e., neurons or muscle). While iron exposure enhanced toxicity resulting from both neuronal and muscle expression of Aβ 42 peptide toxicity at a low temperature, the effects were greater with muscle-specific Aβ 42 peptide expression. Others have also reported differential cell-type Aβ sensitivity and/or toxicity in *C. elegans* [29]. Others assessed Aβ 42 toxicity due to either full-length (Aβ1-42) or truncated (Aβ 3-42) peptide and found greater toxicity associated with the full sequence [69]. Others used the truncated muscle specific expression of Aβ 3-42 peptide with different promoters (*myo-3 vs unc-54*) and noted that *myo-3* had rapid paralysis than *unc-54* promoter [70]. Our study was limited to the truncated Aβ 3-42 peptide, as the truncated Aβ 3-42 is less toxic at low temperature [69, 71]. Regardless, even within the same truncated Aβ 3-42 peptide, we observed cell-type (neuron vs muscle) and promoter (muscle *myo-3* vs *unc-54*) specific responses to iron toxicity at low temperature. Earlier studies reported that divalent metals enhanced Aβ 42 peptide toxicity at high temperature [69, 72]. We observed similar effects at high temperature, but more importantly, iron toxicity exhibited a selective potentiation of Aβ 42 peptide toxicity at low temperature. These results suggest that iron mimics the effects of elevated temperature on Aβ 42 peptide pathology. By extrapolation, iron enriched environments would be expected to activate and enhance otherwise dormant Aβ 42 peptide pathologies. We also found that all treatments were further impacted at higher temperatures, although the degree of pathology was greatest for muscle-specific Aβ 42 peptide expression. Because muscle expressing Aβ 42 peptide also induces a roller phenotype, we ruled out a potential independent role of the roller phenotype by confirming the absence of enhanced iron-induced toxicity in roller worms.

The temperature coefficient (Q_10_) assesses the relative degree of responsiveness of a process to a change in temperature [62]. We tested the hypothesis that Aβ 42 peptide pathology exhibits an increased responsiveness to an elevation in temperature. We confirmed enhanced Aβ 42 peptide pathologies to an increase in temperature according to the following rank order: muscle > neuron > wild type. In contrast, iron enhanced the temperature responsiveness in the following order: wild type > muscle > neuron. The net increase in sensitivity of WT worms following exposure to iron suggests that iron acts synergistically with temperature, while depressing those of muscle- and neuron-specific Aβ 42 peptide expression. Future studies are needed to determine if reduced iron-Aβ 42 peptide synergy is observed at even higher temperatures.

Our data support differential tissue-specific responsiveness of Aβ 42 peptide expression to multiple temperature- and iron-associated pathologies (pumping rate, swimming rate, paralysis). Correlational analyses confirmed that conditions where worms exhibited the lowest pumping and swimming rates also exhibited the highest rates of paralysis, analogous to previous reports in which reduced swimming rate was used as an indication of a decline in energetic function [31]. In this context, our data suggest that iron exposure and Aβ 42 peptide expression may inhibit the activity of key cellular energy producing machinery such as mitochondria. As others have reported, Aβ expressing worms exhibit reduced ATP and oxygen consumption rates [73–75], suggesting that Aβ 42 peptide negatively impacts mitochondria function. Consistent with this idea, we observed reduced mitochondrial bioenergetic function in Aβ 42 peptide expressing worms at baseline, which was exacerbated following iron exposure. Our data suggest that the Aβ effect on mitochondrial bioenergetic function is mediated in part through increased mitochondrial inner membrane permeability or leak. Increased leak results in enhanced ROS production and oxidative damage. Indeed, others have reported that both Aβ 42 peptide and iron cause enhance high levels of ROS and lipid peroxidation by [73–75]. Thus, iron and Aβ 42 may share a similar mechanism of toxicity by promoting mitochondrial bioenergetic dysfunction. When acting together, Aβ 42 and iron would be expected to synergistically exacerbate mitochondrial inner membrane leakiness, bioenergetic dysfunction, ROS production, oxidative stress, and lipid peroxidation. Iron measurements confirmed that while baseline tissue iron levels were minimal and similar for all genotypes, iron exposure resulted in different levels of increased iron burden in WT, roller, neuron- and muscle-specific Aβ 42 expressing worms. Specifically, iron exposure resulted in significantly elevated (>10-fold) iron burden levels in WT, roller, and neuronal Aβ worms, but only a modest elevation (∼2-fold) in muscle-specific Aβ 42 expressing worms. The observation that WT, roller, and neuronal Aβ worms exhibited similar iron burden, but very different levels of toxicity, suggests that iron sensitivity may be increased in neuronal Aβ 42 worms. In line with this idea, iron burden was lowest, and toxicity was highest (reduced pumping and swimming rate, increase paralysis) for muscle-specific Aβ 42 expressing worms. These findings indicate that Aβ 42 peptide expression enhances tissue, and by extension mitochondrial, iron sensitivity, and moreover, that this effect of Aβ 42 peptide is greatest in muscle. This conclusion is supported by a strong correlation for both neuronal and muscle Aβ 42 peptide with respect to each outcome measure.

## Author Contributions

Wilson Peng: Writing – review & editing, Investigation. Kaitlin B. Chung: Writing – review & editing, Investigation. Ali Al-Qazzaz. Writing – review & editing, Investigation, Aidan Straut: Writing – review & editing, Investigation, B. Paige Lawrence: Writing – review & editing, Supervision. M. Kerry O’Banion: Writing – review & editing, Supervision. Robert T. Dirksen: Writing – review & editing, Supervision. John O. Onukwufor: Writing – original draft, Writing – review & editing, Supervision, Methodology, Investigation, Funding acquisition, Formal analysis, Conceptualization.

## Data Availability Statement

The data and all related materials are available upon request.

## Acknowledgments

We thank Drs. Andrew Wojtovich and Keith Nehrke for helpful advice and suggestions. We are grateful to Thomas Scrimale of the Environmental Health Sciences Core (EHSC) and Center for Advanced Research Technologies for elemental analysis. All worm strains used in this study were obtained from the *Caenorhabditis* Genetics Center (CGC), which is supported by NIH Office of Research Infrastructure Programs (P40 OD010440). Graphical image was generated with BioRender. We also appreciate the rigorous discussions and constructive input from the Mitochondrial Research & Innovation Group at the University of Rochester Medical Center, the Western New York Worm Group, and the Pharmacology and Physiology Department Journal Club.

---

**S Fig. 1:**
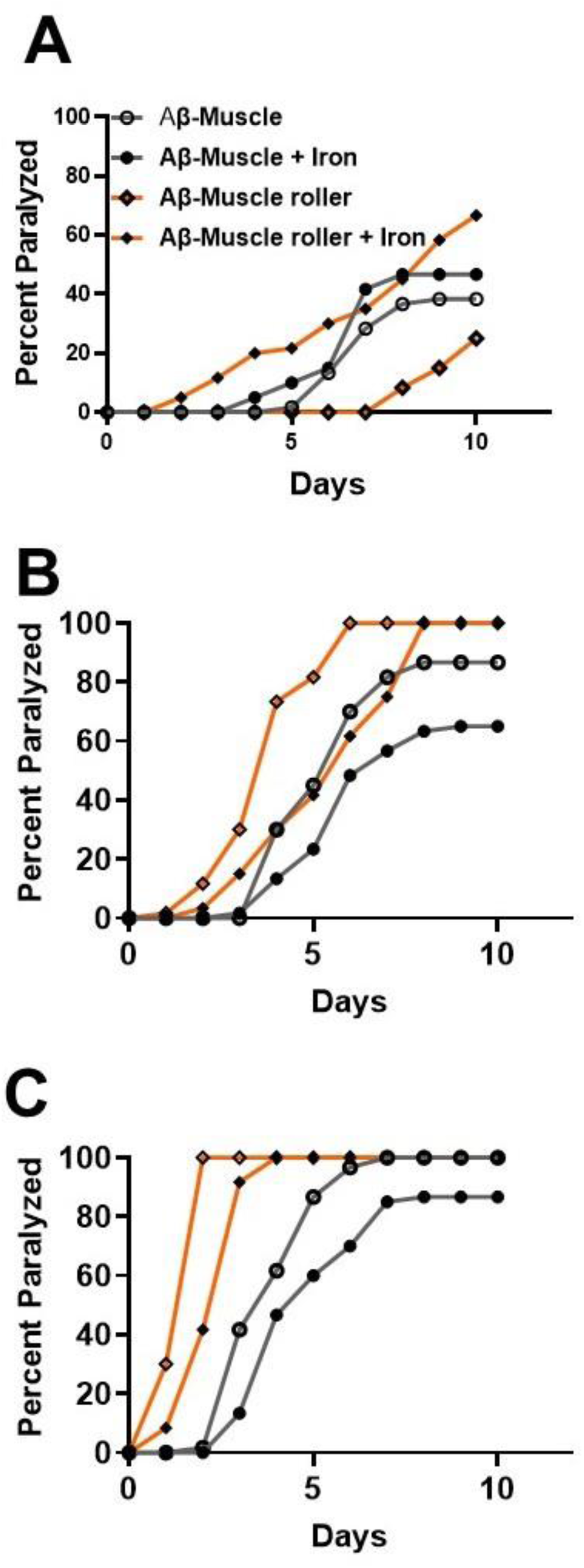
Muscle Aβ 42 peptide thermal and iron sensitivity depends on the promoter. Synchronized L4 worms from Aβ muscle (unc-54) [A-C] were transferred to a plate containing iron (0 and 35 µM) and assigned to temperatures (16, 20 and 25⁰C). Worms were transferred every 24h for 10 days. Worms were scored for paralysis (e.g., inability to move upon stimulation) every 24h for 10 days. **Data from WT Aβ muscle (*myo*-3) was embedded in Aβ muscle (*unc*-54) treatments to see their deviations from Aβ muscle (*myo*-3).** Data are mean of N=3 independent biological replicate (where one biological replicate contains 20 worms per plate).

**Table S1:**
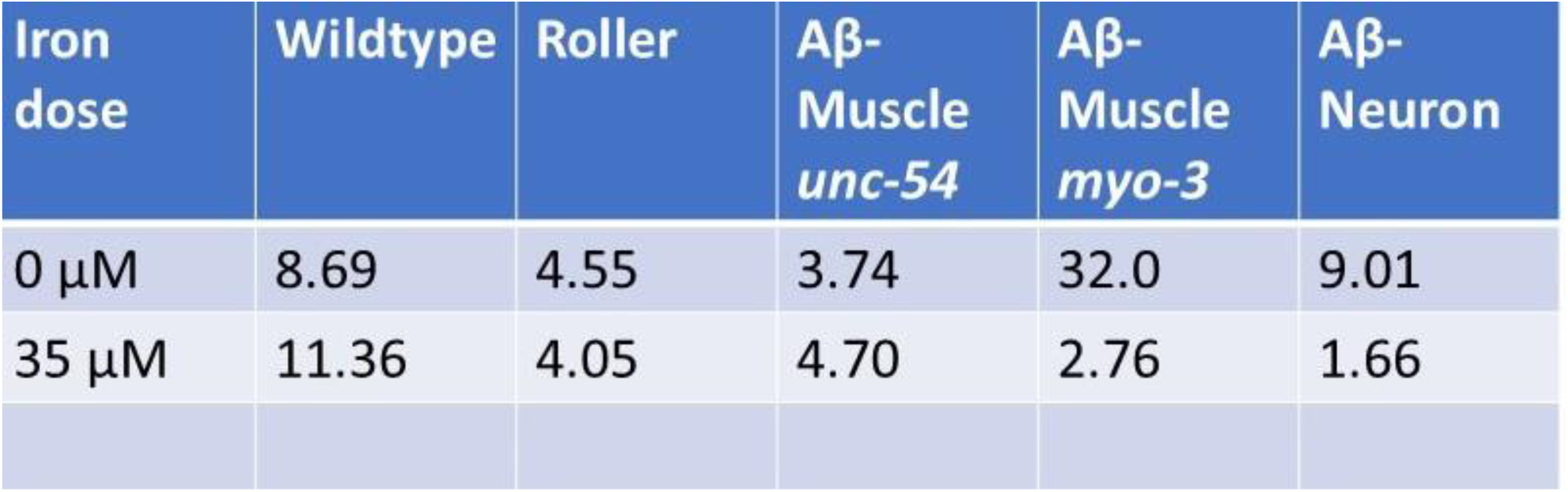
Temperature-dependence of Aβ 42 peptide and iron toxicity. Data are average mean Q 10 values for temperature range 16-20⁰C.

**S Fig. 2:**
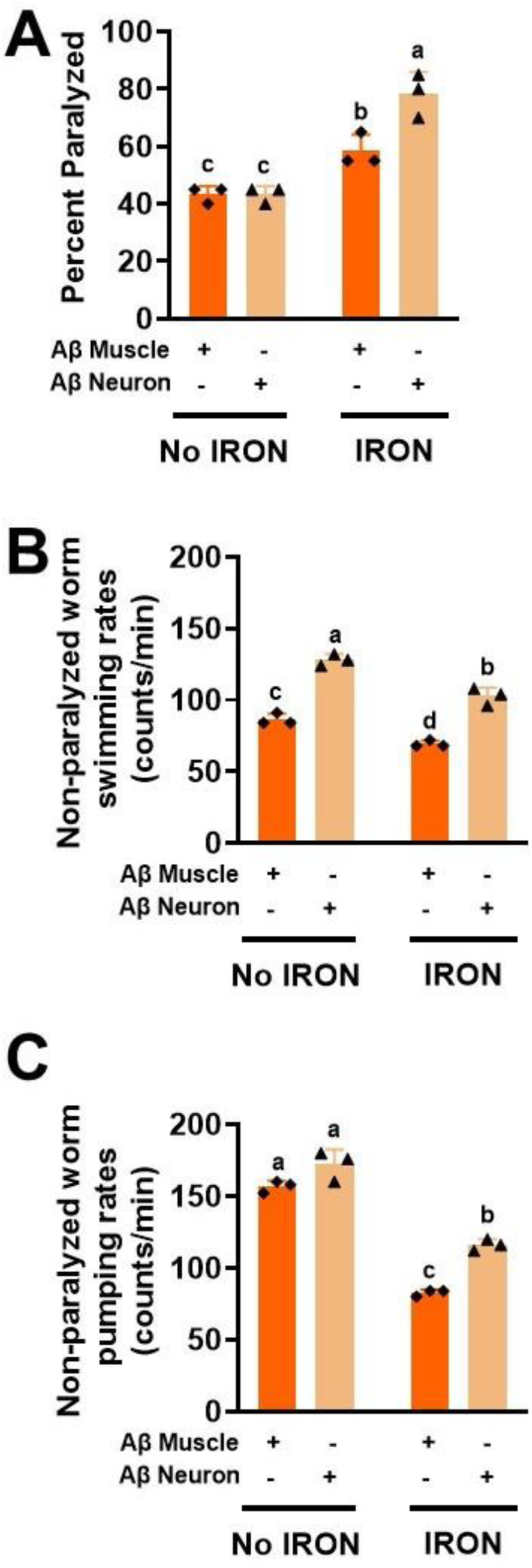
Adjusted Aβ muscle (minus roller) and comparing values with Aβ neuron. A) Percent paralyzed, B) Swimming rate and C) Pumping rate. Data are mean ± SEM, N = 3 independent replicates (where 5 independent worm count constitute an N). Bars with different letter differ significantly, two-way ANOVA, Tukey post hoc test.

