## Supplemental Data 1 for "Iron toxicity potentiates cell-type specific amyloid beta proteotoxicity in *C. elegans* via altered energy homeostasis"

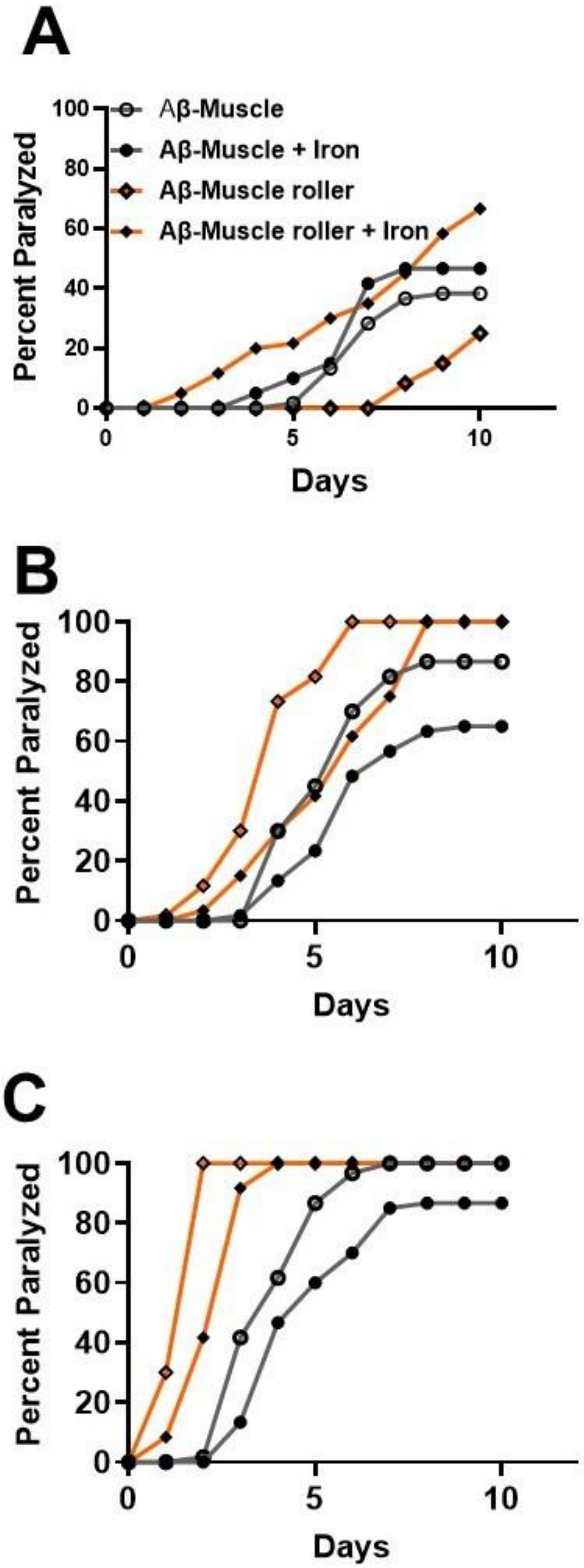

A
Aβ-Muscle
Aβ-Muscle + Iron
Aβ-Muscle roller
Aβ-Muscle roller + Iron
Percent Paralyzed
Days
B
Percent Paralyzed
Days
C
Percent Paralyzed
Days

**S Fig. 1: Muscle Aβ 42 peptide thermal and iron sensitivity depends on the promoter**. Synchronized L4 worms from Aβ muscle (unc-54) [A-C] were transferred to a plate containing iron (0 and 35 μM) and assigned to temperatures (16, 20 and 25ºC). Worms were transferred every 24h for 10 days. Worms were scored for paralysis (e.g., inability to move upon stimulation) every 24h for 10 days. **Data from WT Aβ muscle (*myo*-3) was embedded in Aβ muscle (*unc*-54) treatments to see their deviations from Aβ muscle (*myo*-3).** Data are mean of N=3 independent biological replicate (where one biological replicate contains 20 worms per plate).

S Table 1: Temperature-dependence of Aβ 42 peptide and iron toxicity. Data are average mean Q 10 values for temperature range 16-20ºC.

| Iron dose | Wildtype | Roller | Aβ-Muscle *unc-54* | Aβ-Muscle *myo-3* | Aβ-Neuron |
|---|---|---|---|---|---|
| 0 µM | 8.69 | 4.55 | 3.74 | 32.0 | 9.01 |
| 35 µM | 11.36 | 4.05 | 4.70 | 2.76 | 1.66 |

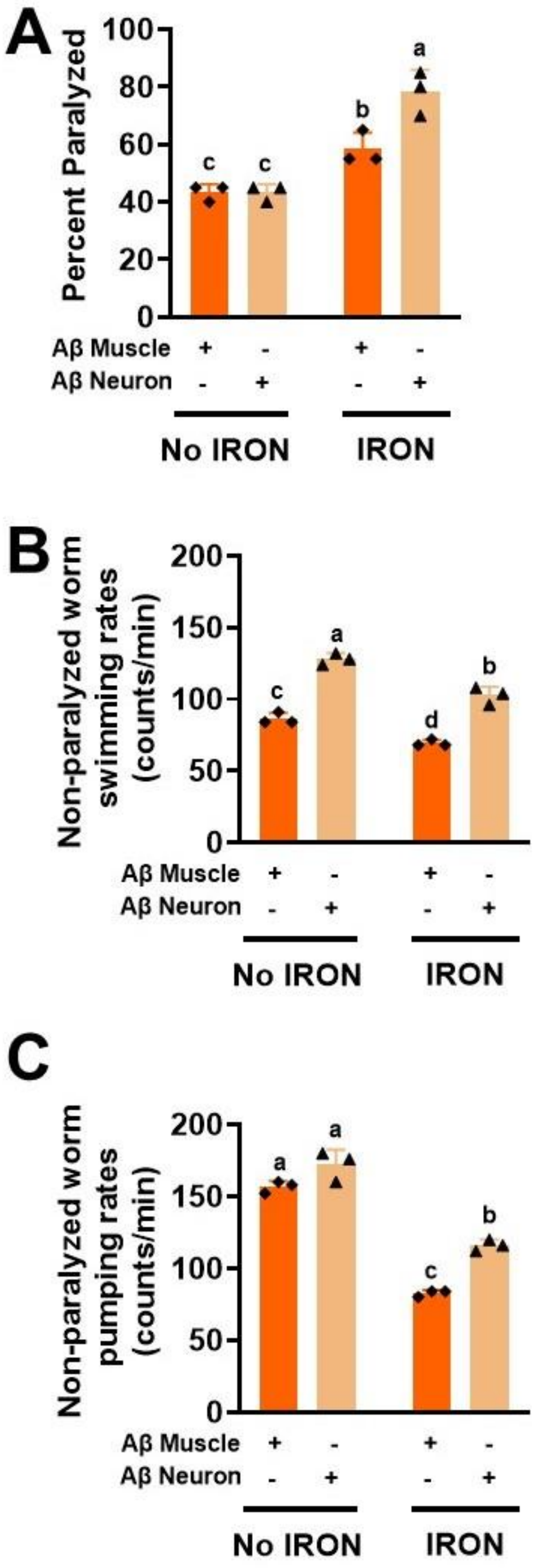
A
Percent Paralyzed
100
80
60
40
20
0
c
c
b
a
Aβ Muscle + - + -
Aβ Neuron - + - +
No IRON
IRON
B
Non-paralyzed worm swimming rates (counts/min)
200
150
100
50
0
c
a
d
b
Aβ Muscle + - + -
Aβ Neuron - + - +
No IRON
IRON
C
Non-paralyzed worm pumping rates (counts/min)
200
150
100
50
0
a
a
c
b
Aβ Muscle + - + -
Aβ Neuron - + - +
No IRON
IRON

**S Fig. 2: Adjusted Aβ muscle (minus roller) and comparing values with Aβ neuron**. A) Percent paralyzed, B) Swimming rate and C) Pumping rate. Data are mean ± SEM, N = 3 independent replicates (where 5 independent worm count constitute an N). Bars with different letter differ significantly, two-way ANOVA, Tukey post hoc test.
